# Decoding the absolute stoichiometric composition and structural plasticity of α-carboxysomes

**DOI:** 10.1101/2021.12.06.471529

**Authors:** Yaqi Sun, Victoria M. Harman, James R. Johnson, Taiyu Chen, Gregory F. Dykes, Yongjun Lin, Robert J. Beynon, Lu-Ning Liu

## Abstract

Carboxysomes are anabolic bacterial microcompartments that play an essential role in carbon fixation in cyanobacteria and some chemoautotrophs. This self-assembling organelle encapsulates the key CO_2_-fixing enzymes, Rubisco, and carbonic anhydrase using a polyhedral protein shell that is constructed by hundreds of shell protein paralogs. The α-carboxysome from the chemoautotroph *Halothiobacillus neapolitanus* serves as a model system in fundamental studies and synthetic engineering of carboxysomes. Here we adopt a QconCAT-based quantitative mass spectrometry to determine the absolute stoichiometric composition of native α-carboxysomes from *H. neapolitanus*. We further performed an in-depth comparison of the protein stoichiometry of native and recombinant α-carboxysomes heterologously generated in *Escherichia coli* to evaluate the structural variability and remodeling of α-carboxysomes. Our results provide insight into the molecular principles that mediate carboxysome assembly, which may aid in rational design and reprogramming of carboxysomes in new contexts for biotechnological applications.

## Introduction

Bacterial microcompartments (BMCs) are self-assembling proteinaceous organelles that are widespread among bacterial phyla (Axen et al., 2014; Sutter et al., 2021). The BMC is composed of a virus-like polyhedral protein shell that sequesters a series of enzymes to segregate their metabolic processes from the cytoplasm and provide specific local microenvironments to favor enzymatic activities (Kerfeld et al., 2018; Liu, 2021a; Liu, 2021b; Yeates et al., 2008). Increasing evidence has been achieved to highlight the significant roles of BMCs in enhancing the metabolism of various carbon sources, alleviating metabolic crosstalk, and encapsulating toxic/volatile metabolites (Bobik et al., 2015; Chowdhury et al., 2014; Greening and Lithgow, 2020).

Carboxysomes are anabolic BMCs for autotrophic CO_2_ fixation in all identified cyanobacteria and some chemoautotrophs (Borden and Savage, 2021; Kerfeld et al., 2018; Liu, 2021a; Rae et al., 2013; Sun et al., 2020). They encase the CO_2_-fixing enzymes, Ribulose-1,5-bisphosphate carboxylase oxygenase (Rubisco) and carbonic anhydrase (CA), using a semi-permeable shell, which allows the passage of negatively charged HCO_3-_ and Ribulose 1,5-bisphosphate (RuBP) and probably preclude O_2_ influx and leakage of CO_2_ from the carboxysome to the cytoplasm (Dou et al., 2008; Faulkner et al., 2020; Mahinthichaichan et al., 2018). In the carboxysome lumen, HCO_3-_ is dehydrated to CO_2_ by CA, ensuring elevated CO_2_ levels around Rubisco to facilitate Rubisco carboxylation and reduce wasteful photorespiration (Long et al., 2021; Price et al., 2008). Collectively, the intriguing self-assembly and selective permeability features of carboxysomes provide the structural basis for enhanced CO_2_ assimilation and substantial contributions to global primary production (Hennacy and Jonikas, 2020; Rae et al., 2013; Rae et al., 2017).

According to the forms of encapsulated Rubisco and protein composition, carboxysomes can be categorized into two sub-classes, α- and β-carboxysomes (Kerfeld and Melnicki, 2016; Rae et al., 2013). The α-carboxysome of the chemoautotrophic bacterium *Halothiobacillus neapolitanus* (hereafter as *H. neapolitanus*) has been chosen as a model carboxysome in fundamental studies and synthetic engineering. The genes encoding α-carboxysome-related proteins are clustered mainly in the *cso* operon in the *H. neapolitanus* genome (Figure 1). The shell is constructed by six types of paralogous proteins, including the hexameric proteins (BMC-H) CsoS1A, CsoS1B and CsoS1C that tile the major facet of shells, the pentamers (BMC-P) CsoS4A and CsoS4B that sit at the vertexes, and the trimeric pseudo-hexamer (BMC-T) CsoS1D that possesses a larger central pore than other shell proteins, which was proposed to play a role in mediating the passage of large metabolite molecules, such as RuBP and 3-phosphoglycerate (3-PGA) (Bonacci et al., 2012; Faulkner et al., 2020; Klein et al., 2009; Roberts et al., 2012). Among the BMC-H proteins, CsoS1A and CsoS1C have a high sequence similarity, differing in only 2 amino acids out of 98 (Heinhorst and Cannon, 2020; Tsai et al., 2007), whereas CsoS1B contains a 12-residue C-terminal extension (Tsai et al., 2007). The cargo enzymes include Rubisco and CA. Rubisco is assembled by the large and small subunits CbbL and CbbS that form an L_8_S_8_ hexadecamer. CsoSCA acts as the functional CA in the α-carboxysome, existing as a dimer (Sawaya et al., 2006), and was suggested to associate with the shell inner surface (Cai et al., 2015; Dou et al., 2008). The linker protein CsoS2 in the *H. neapolitanus* α-carboxysome has two isoforms, a shorter polypeptide CsoS2A (C-terminus truncated) and a full-length CsoS2B, translated via programmed ribosomal frame shifting (Chaijarasphong et al., 2016). CsoS2A and CsoS2B shared the middle region and the N-terminal domain that binds with Rubisco and induces Rubisco condensation (Oltrogge et al., 2020). The C-terminus of CsoS2B, which is absent in CsoS2A, is presumed to bind with the shell and can serve as an encapsulation peptide to recruit non-native cargos (Cai et al., 2015; Li et al., 2020). In addition, CbbO and CbbQ function as the Rubisco activases, forming a bipartite complex comprising one CbbQ hexamer and one CbbO monomer, to remove inhibitors from the Rubisco catalytic site to restore its carboxylation (Chen et al., 2021; Sutter et al., 2015; Tsai et al., 2015; Tsai et al., 2020).

**Figure 1.**
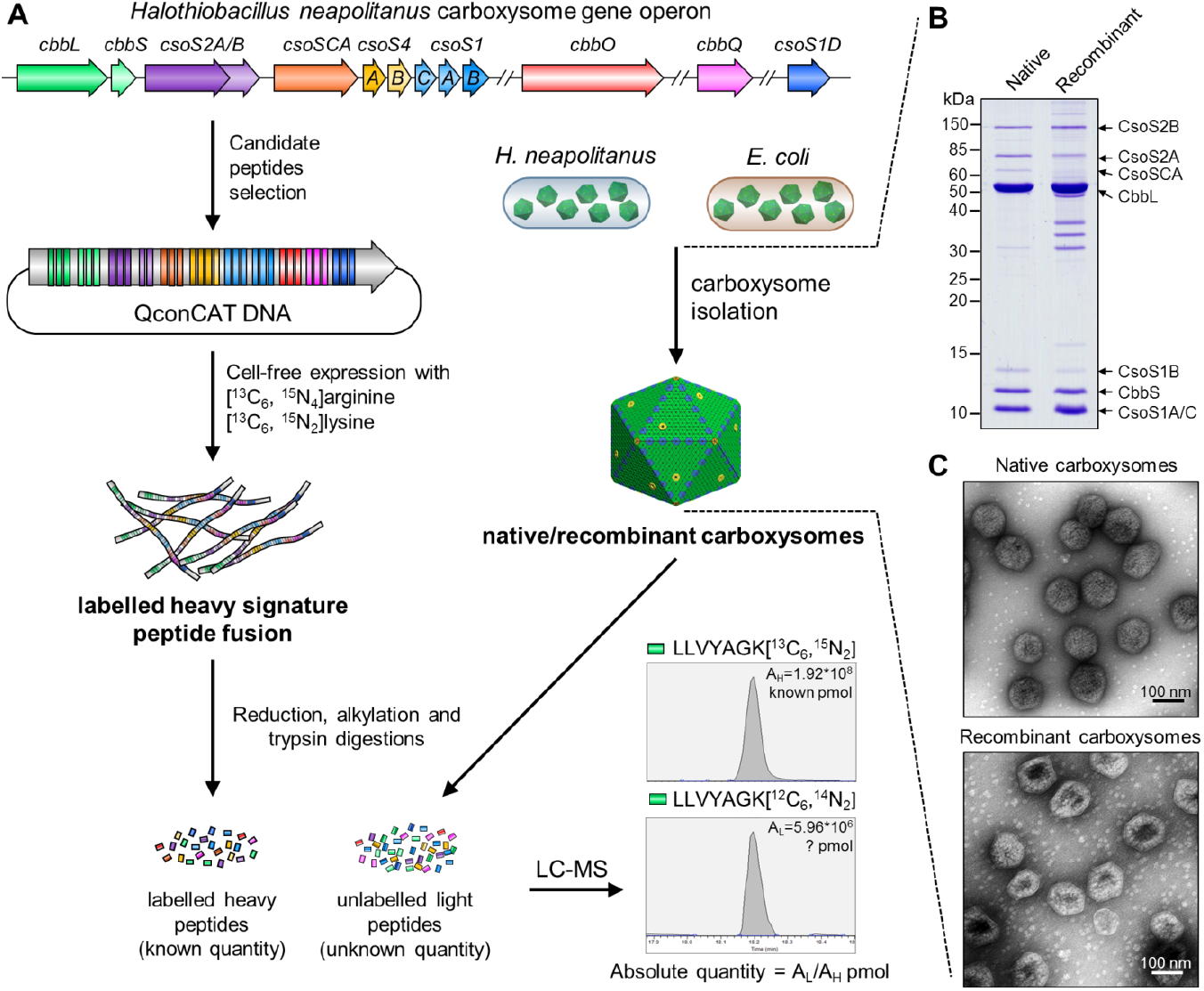
Schematic overview of QconCAT strategy. (**A**) QconCAT DNA fragment was designed from selected gene sequences from the *H. neapolitanus* operon that expresses α-carboxysome proteins. The stable isotopes ([^13^C_6_,^15^N_4_] arginine and [^13^C_6_, ^15^N_2_] lysine) labelled QconCAT peptide fusion was expressed via a cell-free system, purified and quantified and added to four replicates samples of isolated native/recombinant α-carboxysomes from *H. neapolitanus* and *E. coli*. The absolute quantity and stoichiometry of unlabelled signature peptides for carboxysomal proteins were calculated by LC-MS analysis. One signature peptide for CbbQ, LLVKAGK was shown here as an example. (**B**) SDS-PAGE of isolated native/recombinant α-carboxysomes showing the majority bands of α-carboxysome proteins. (**C**) EM images of isolated native/recombinant α-carboxysomes.

Given the significance of metabolic improvement and synthetic engineering potential, substantial efforts have been made to uncover the assembly and structural principles of carboxysomes. However, our knowledge about the accurate stoichiometric composition of carboxysomes, which plays an essential role in determining their size, shape, structural integrity, permeability, and catalytic performance (Liu et al., 2021), is still primitive. Label-free quantitative mass spectrometry has been used to determine the relative content of protein compositions within the BMCs (Faulkner et al., 2017; Havemann and Bobik, 2003; Long et al., 2005; Mayer et al., 2016). Furthermore, our recent work has applied mass spectrometry-based absolute quantification and a QconCAT (concatamer of standard peptides for absolute quantification) strategy to examine the precise stoichiometric composition of 1,2-propanediol utilization (PDU) metabolosomes from *Salmonella enterica* serovar Typhimurium LT2 (Yang et al., 2020). In addition, fluorescence labeling and microscopic imaging have been utilized to characterize the protein stoichiometry of β-carboxysomes from the cyanobacterium *Synechococcus elongatus* PCC 7942 (Syn7942) (Sun et al., 2019). However, the precise stoichiometric composition of α-carboxysomes has not been well characterized, despite the crude estimates based on protein electrophoresis profiles reported in previous studies (Cannon and Shively, 1983; Heinhorst et al., 2006a; Roberts et al., 2012).

Here, we perform absolute quantification of protein components within native α-carboxysomes from *H. neapolitanus* and recombinant α-carboxysomes produced in *Escherichia coli* (*E. coli*), using QconCAT-assisted quantitative mass spectrometry in combination with biochemical analysis, electron microscopy (EM) and enzymatic assays. Our results shed light on the molecular principles underlying the assembly and structural plasticity of α-carboxysomes, and provide essential information required for design and engineering of carboxysomes in synthetic biology.

## Results

### Quantifying the protein stoichiometry of native α-carboxysomes from *H. neapolitanus*

The QconCAT-assisted mass spectrometry approach permitted a precise quantification of the absolute abundance of proteins (Johnson et al., 2021; Rivers et al., 2007; Simpson and Beynon, 2012). This approach has been recently applied to quantify the stoichiometric composition of protein components within the Pdu metabolosome (Yang et al., 2020). To determine the stoichiometry of α-carboxysome components, native α-carboxysomes were first isolated from *H. neapolitanus* using sucrose gradient ultracentrifugation (Figure S1A). Sodium dodecyl sulfate–polyacrylamide gel electrophoresis (SDS-PAGE) indicated that CsoS2A/B, CbbL/S, and CsoS1A/B/C are the major α-carboxysomes proteins (Figure S1B). NADH-coupled CO_2_-fixation activity assays confirmed the functionality of isolated α-carboxysomes, with a measured carbon fixation *V*_*max*_ of 2.96 ± 0.09 μmol.mg^-1.^min^-1^, and *K*_*m*(RuBP)_ at 0.20 ± 0.02 mM (*n* = 4) (Figure S1C). EM showed that the isolated α-carboxysomes form intact and canonical polyhedral shape, with an average diameter of 124.6 ± 9.6 nm (*n* = 272) (Figure S1D, S1E), consistent with previous results (Holthuijzen et al., 1986; Shively et al., 1973; Sutter et al., 2015).

To establish the accurate stoichiometry of all proteins within the isolated α-carboxysomes, we used high-resolution liquid chromatography-mass spectrometry (LC-MS) calibrated with protein-specific stable-isotope labeled internal standards generated via the QconCAT strategy (Figure 1, Figure S2) (Johnson et al., 2021; Pratt et al., 2006). The designed single QconCAT peptide is composed of 3 unique peptides for CbbL, CbbS, CsoSCA, CbbO, CbbQ, CsoS1D, CsoS4A and CsoS2AB shared region, 2 peptides for CsoS2B and CsoS1ABC shared region, as well as 1 peptide for the CsoS1B and CsoS1AC shared region (Figure S2A, Table S1). Due to the high sequence similarity, CsoS1A and CsoS1C could not be distinguished in the current QconCAT design. The QconCAT peptide also contains peptides of the form II Rubisco CbbM. Since CbbM was not presumed to be a component of the *H. neapolitanus* α-carboxysome (Baker et al., 1998), we used CbbM as a reference to validate the quality of α-carboxysome isolation. The genes encoding these selected peptide candidates were assembled, following the Qbrick assembly strategy (Johnson et al., 2021), to form the QconCAT DNA sequence (Table S2). The designed QconCAT peptide was then produced by cell-free synthesis (Takemori et al., 2017) and was further isolated, validated by SDS-PAGE (Figure S2B).

MS1 precursor QconCAT quantification was then carried out using four batches of independently isolated α-carboxysomes (Figure 1A, Figure S3). The purified α-carboxysomes were mixed with the QconCAT standard, co-digested, and analyzed by label-free MS quantification. All carboxysomal proteins were detected in the isolated carboxysomes, whereas CbbM was not detectable in the isolated samples. The carboxysomal proteins account for 99.5 ± 0.2% of the total proteins in the samples, confirming the high purity of isolated carboxysomes (Figure S4A). Accuracy and reliability of protein quantification were verified by a good agreement of the peptides for each carboxysome protein in the four biological replicates (Figure S4C).

We quantified the abundance of protein components within one *H. neapolitanus* carboxysome structure, based on the shell surface area of a typical icosahedron (Whitehead et al., 2014) and the average carboxysome size (124.6 ± 9.6 nm, *n* = 272) measured in EM (Figure 1C, Table 1, Table S3, see details in Methods). The results revealed that the most abundant proteins in the *H. neapolitanus* α-carboxysome are CsoS1AC hexamers (863 copies), followed by Rubisco (447 copies, estimated by the CbbL content), CsoS2A (248 copies), CsoS2B (192 copies), CsoS1B hexamers (112 copies), and 58 copies of CsoSCA dimers. The *H. neapolitanus* α-carboxysome has a molecular weight (MW) of ∼346 MDa and the Rubisco enzymes account for ∼66% of the total MW. The hexameric shell proteins CsoS1A/C and CsoS1B make up ∼17.1% of the total MW. Additionally, 11 copies of CsoS4A/B pentamers (CsoS4A: 8.8; CsoS4B: 2.2) are integrated within the α-carboxysome, slightly less than 12 that is typically assumed to cap the vertices of an icosahedron. CsoS1D pseudo-hexamers have a low abundance in the shell, with 2.9 copies per carboxysome. Moreover, the linker proteins, CsoS2A and CsoS2B, account for 13.5% of the total MW.

**Table 1.** QconCAT quantification of protein components in native α-carboxysomes from *H. neapolitanus*.

| Category | Protein | Structure of functional unit | MW (kDa) | % Total protein by weight | Unit of monomeric protein per carboxysome | Functional unit of multimer per carboxysome |
| --- | --- | --- | --- | --- | --- | --- |
| Structural Proteins | CsoS1AC | Hexamer (Tsai et al., 2007; Tsai et al., 2009) | 10.0 | 14.88 ± 1.09 | 5175.1 ± 378.1 | 862.5 ± 63 |
| | CsoS1B | Hexamer* | 11.3 | 2.20 ± 0.48 | 673.3 ± 148.3 | 112.2 ± 24.7 |
| | CsoS1D | Pseudo-hexamer (Klein et al., 2009) | 23.5 | 0.12 ± 0.01 | 17.4 ± 1.0 | 2.9 ± 0.2 |
| | CsoS4A | Pentamer (Tanaka et al., 2008) | 8.9 | 0.11 ± 0.00 | 43.9 ± 1.6 | 8.8 ± 0.3 |
| | CsoS4B | Pentamer (Zhao et al., 2019) | 8.8 | 0.03 ± 0.00 | 10.8 ± 1.6 | 2.2 ± 0.3 |
| | CsoS2A | Monomer (Cai et al., 2015; Oltrogge et al., 2020) | 92.4 | 6.61 ± 1.26 | 247.7 ± 47.3 | 247.7 ± 47.3 |
| | CsoS2B | Monomer (Cai et al., 2015; Oltrogge et al., 2020) | 124.2 | 6.87 ± 0.54 | 191.5 ± 15.0 | 191.5 ± 15.0 |
| Catalytic proteins | CbbL | L<sub>8</sub>S<sub>8</sub> Hexadecamer (Oltrogge et al., 2020) | 52.6 | 54.36 ± 2.30 | 3575.7 ± 151.3 | 447.0 ± 18.9 |
| | CbbS | L<sub>8</sub>S<sub>8</sub> Hexadecamer (Oltrogge et al., 2020) | 12.9 | 11.74 ± 0.56 | 3161.2 ± 150.7 | 395.1 ± 18.8 |
| | CsoSCA | Dimer (Sawaya et al., 2006) | 57.3 | 1.93 ± 0.12 | 116.6 ± 7.1 | 58.3 ± 3.6 |
| | CbbQ | Hexamer (Sutter et al., 2015) | 30.1 | 0.76 ± 0.05 | 87.8 ± 5.3 | 14.6 ± 0.9 |
| | CbbO | Monomer (Sutter et al., 2015; Tsai et al., 2020) | 88.6 | 0.39 ± 0.03 | 15.4 ± 1.3 | 15.4 ± 1.3 |
| Intact native α-carboxysome | | | 346.3 | | | |

\*Based on the structural similarity with CsoS1A/C.

Approximately 15 copies of CbbQO complexes, each composed of one CbbQ hexamer and one CbbO monomer, were identified in the carboxysome, indicating that the CbbQO complex is a structural component of native α-carboxysomes in *H. neapolitanus*. Consistently, CbbQ has been indicated to be tightly associated with the *H. neapolitanus* carboxysome shell (Sutter et al., 2015) and CbbQO can be incorporated into recombinant α-carboxysomes (Chen et al., 2021). Likewise, our mass spectrometry results showed the presence of McdAB-like proteins in purified native α-carboxysomes (Figure S4A, Supplemental File 1), implicating the association of McdAB-like proteins with α-carboxysomes, which was proposed to ensure proper distribution of α-carboxysomes in *H. neapolitanus* and carboxysome inheritance during cell division (MacCready et al., 2021). Some chemoautotrophs, including *H. neapolitanus*, contain the *cbbM* gene encoding Form II Rubisco and its activases CbbQ1 and CbbO1 (Tsai et al., 2015). These proteins were not detected in the purified carboxysomes (Supplemental File 1), suggesting that they are not the organizational components of or associated with the α-carboxysomes in *H. neapolitanus*.

### Stoichiometric composition of recombinant α-carboxysomes

Previous studies have demonstrated that heterologous engineering of the *H. neapolitanus* α-carboxysomes could result in functional α-carboxysome structures (Baumgart et al., 2017; Bonacci et al., 2012; Chen et al., 2021; Flamholz et al., 2020). To verify the compositional similarity between native and recombinant α-carboxysomes, we reconstituted *H. neapolitanus* α-carboxysomes by expressing the *cso* operon with *csoS1D* using an arabinose-inducible pBAD33 vector in *E. coli* (Figure S1G). SDS-PAGE revealed an overall similar content of protein components within the isolated native and recombinant α-carboxysomes, except for a reduction in the CsoSCA content in recombinant carboxysomes (Figure S1B, S3). Carbon-fixation kinetics as a function of RuBP concentrations confirmed the function of recombinant α-carboxysomes, with a *V*_*max*_ of 2.07 ± 0.12 μmol·mg^-1^·min^-1^ (*n* = 4) and a *K*_*m*(RuBP)_ of 0.08 ± 0.02 mM (*n* = 4), although both were lower than those of native α-carboxysomes (Figure S1C). EM indicated that recombinant α-carboxysomes possess a polyhedral shape and an average diameter of 131.8 ± 18.0 nm (*n* = 152), slightly larger than native α-carboxysomes (Figure S1D, S1E). Analysis of EM images showed that both native and recombinant α-carboxysomes possess single-layer shells (5.3 ± 0.6 nm and 5.5 ± 0.8 nm, respectively, *n* = 100, Figure S1F), consistent with previous observations (Faulkner et al., 2017).

Isolated recombinant α-carboxysomes were then subject to MS1 precursor QconCAT quantification and normalization to retrieve the stoichiometric content of a single carboxysome (Figure 1, Table 2, Table S3). Within the recombinant α-carboxysome, the most abundant proteins are CsoS1AC hexamers (1001 copies), followed by Rubisco (426 copies), CsoS2A (305 copies), CsoS2B (249 copies), and CsoS1B hexamers (79 copies). The recombinant α-carboxysome has a molecular mass of ∼336 MDa, and has significantly reduced Rubisco copy numbers compared with the native α-carboxysome (*p* < 0.05, Figure 2). The content of CsoSCA in the recombinant α-carboxysome is reduced by 29-fold compared to that in the native α-carboxysome, resulting in only ∼2 CsoSCA dimers per recombinant α-carboxysome, consistent with SDS-PAGE analysis (Figure S1B). The hexameric shell proteins, CsoS1AC and CsoS1B, account for 19.4% of the total MW in recombinant α-carboxysomes (Table 2). The CsoS1B content is reduced by ∼30% (79 copies) compared to that in native α-carboxysomes (112 copies, *p* < 0.05, Figure 2). There are on average 7.1 copies of pentameric proteins (CsoS4A: 6.3; CsoS4B: 0.8) in recombinant α-carboxysomes, less than the hypothetical 12 pentamers for a typical icosahedral structure. It suggests that some vertices are not capped by CsoS4 pentamers. Similar features have also been observed in β-carboxysomes and synthetic BMC shells (Hagen et al., 2018; Sun et al., 2019; Sutter et al., 2019), presumably providing a mechanism for regulating shell architecture and permeability. CsoS1D has ∼0.8 copies per recombinant α-carboxysome, less than that in the native α-carboxysome (*p* < 0.001, Table 2). CsoS2A and CsoS2B have 305 and 249 copies, respectively, per recombinant α-carboxysome, collectively accounting for 17.6% of the total MW. CsoS2B has an increased content in the recombinant α-carboxysome than in the native form (Figure 2).

**Table 2.** QconCAT quantification of protein components in recombinant *H. neapolitanus* α-carboxysomes.

| Category | Protein | MW<br>(kDa) | % Total protein<br>by weight | Unit of monomeric<br>protein per<br>carboxysome | Functional Unit<br>of multimer per<br>carboxysome |
| --- | --- | --- | --- | --- | --- |
| Structural<br>Proteins | CsoS1AC | 10.0 | 17.80 ± 0.80 | 6002.9 ± 268.1 | 1000.5 ± 44.7 |
| | CsoS1B | 11.3 | 1.59 ± 0.19 | 473.1 ± 56.6 | 78.8 ± 9.4 |
| | CsoS1D | 23.5 | 0.03 ± 0.00 | 5.0 ± 0.3 | 0.8 ± 0.1 |
| | CsoS4A | 8.9 | 0.08 ± 0.01 | 31.7 ± 3.5 | 6.3 ± 0.7 |
| | CsoS4B | 8.8 | 0.01 ± 0.01 | 3.9 ± 2.7 | 0.8 ± 0.5 |
| | CsoS2A | 92.4 | 8.39 ± 0.26 | 305.0 ± 9.3 | 305.0 ± 9.3 |
| | CsoS2B | 124.2 | 9.21 ± 0.47 | 249.0 ± 12.6 | 249.0 ± 12.6 |
| Catalytic<br>proteins | CbbL | 52.6 | 53.42 ± 3.96 | 3408.1 ± 252.4 | 426.0 ± 31.5 |
| | CbbS | 12.9 | 9.38 ± 0.37 | 2449.4 ± 96.2 | 306.2 ± 12 |
| | CsoSCA | 57.3 | 0.07 ± 0.02 | 4.0 ± 1.1 | 2.0 ± 0.5 |
| **Intact recombinant α-carboxysome** |  | **335.8** |  |  |  |

**Figure 2.**
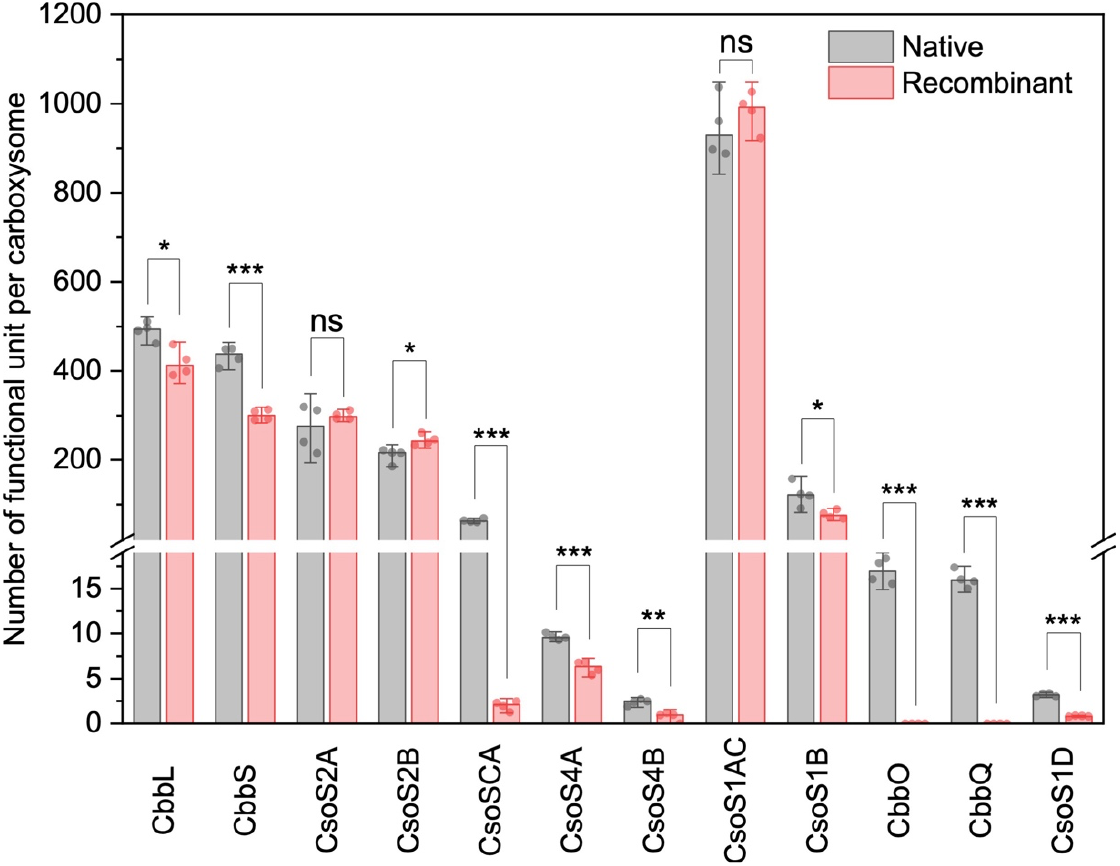
Stoichiometry comparison of native and recombinant carboxysomes. ns, as no statistical significance; **\***, *p* < 0.05; **\*\***, *p* < 0.01; **\*\*\***, *p* < 0.001 using two sample t-test, equal variance not assumed (welch correction). Data are shown as mean ± SD from four biological replicates.

## Discussion

In this study, we performed absolute quantification using QconCAT-based mass spectrometry to determine the stoichiometric composition of the *H. neapolitanus* α-carboxysomes, which represent a step toward gaining a comprehensive understanding of the structure and function of the model carboxysome.

Given that BMC components have a notable variation in protein abundance (Yang et al., 2020) and some minor proteins were not identifiable as well as the protein paralogs with similar molecular were not distinguishable in SDS-PAGE gels (Figure 1, Figure S2), it is difficult to obtain the accurate protein stoichiometry of carboxysomes based on protein electrophoresis profiles. Comparison of QconCAT and label-free quantification results illustrated marked deviations in the abundance of some carboxysomal proteins (Figure S4B). The results demonstrated that label-free quantification could potentially underestimate the content of CsoS1B, CsoSCA, CsoS4A, and CsoS4B by 48/32%, 64/95%, 144/142%, and 119/105% (native/recombinant carboxysomes), respectively, highlighting the necessity of QconCAT-based quantification in studying the protein stoichiometric composition of BMCs.

### Stoichiometric variability and structural plasticity of α-carboxysomes

Characterization of the absolute stoichiometric compositions for native and recombinant carboxysomes provides insight into the organizational principles and plasticity of the *H. neapolitanus* α-carboxysome (Figure 3). It becomes apparent that the BMC shells are amendable to integrate different copies or types of shell proteins, and the absence of specific components or the changes in the ratios of protein paralogs may not necessarily impede the overall shell assembly (Garcia-Alles et al., 2019; Long et al., 2018; Sommer et al., 2019; Yang et al., 2020). The total copy number of shell pentamers (CsoS4A and CsoS4B) is 11.0 for native α-carboxysomes and 7.1 for recombinant α-carboxysomes, both less than 12 pentamers that are postulated to occupy all the vertices of a regular icosahedron (Bobik et al., 2015; Kerfeld et al., 2018). These results elucidated that it is not a prerequisite to cap all the vertices with pentamers in a functional carboxysome. In support of this, polyhedral carboxysomes and BMC shells deficient in pentamers could still be formed (Cai et al., 2009; Hagen et al., 2018; Lassila et al., 2014; Long et al., 2018). Our previous study has also demonstrated that variable copies of CcmL pentamers are integrated in Syn7942 β-carboxysomes under different growth conditions (Sun et al., 2019). The lack of pentamers at some vertices might result in observable structural heterogeneity and reduced integrity of the entire α-carboxysomes (Figure S1D).

**Figure 3.**
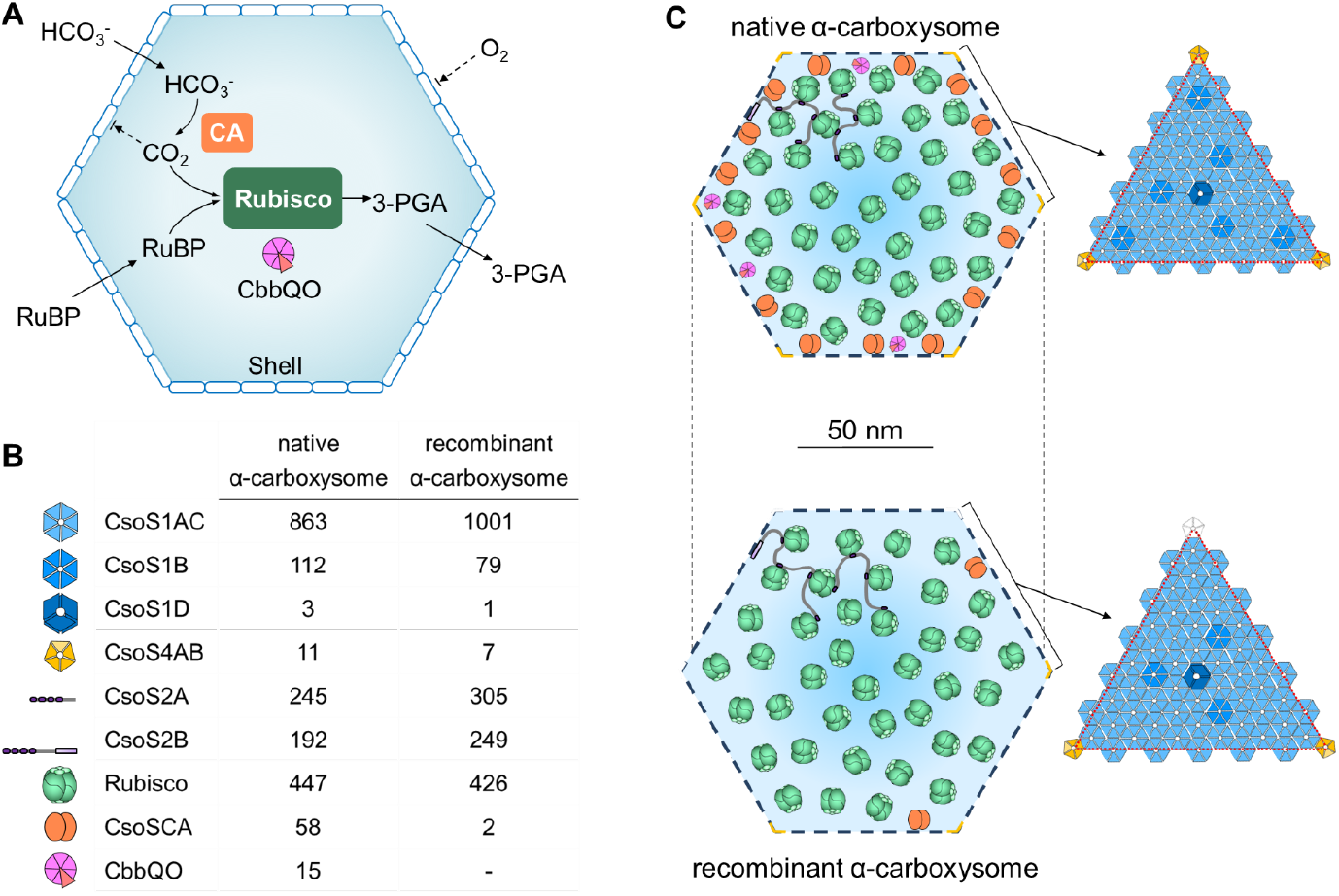
Structural models of *H. neapolitanus* α-carboxysomes. (**A**) Schematic of the pathways of carbon fixation in the α-carboxysome, including Rubisco activases CbbQO as the structural components; (**B**) The stoichiometry of each structural component within native and recombinant α-carboxysome (see Table 1 and 2). (**C**) Schematic of native and recombinant α-carboxysome structures and shell organizations. The numbers of proteins do not represent actual abundance and is only for illustration purposes.

Rubisco in carboxysomes was proposed to adopt a Kepler packing, filling maximally 74% of the internal carboxysome volume (Long et al., 2011; Whitehead et al., 2014). Quantification based upon the CbbL content indicates that the native *H. neapolitanus* α-carboxysome can accommodate approximately 447 Rubisco (the CbbL:CbbS ratio is 8:7.3), in agreement with the theoretical estimation based on the Kepler packing (411 Rubisco, Table S3). In contrast, recombinant α-carboxysomes encapsulate 426 Rubisco (the CbbL:CbbS ratio is 8:5.7), lower than the estimated copy number of 491 based on measured recombinant carboxysome size (Table S3). The increased shell:interior ratio (from 0.8:1 to 1:1, Table. 3) and carboxysome size specified a lower packing density of Rubisco within recombinant carboxysomes (Figure 3). Moreover, the perturbed formation of Rubisco (L_8_S_8_) as indicated by the changes in the CbbL:CbbS ratio has also been determined in recombinant carboxysomes (Table 3). Our results also showed that the Rubisco/CA (CbbL:CsoSCA) ratio varies drastically between native and recombinant α-carboxysomes (Table 3). It has been postulated that too little or too much carboxysomal CA activity, which could cause limited CO_2_ supply or substantial leakage of CO_2_, may interfere with CO_2_ fixation of carboxysomes (Rae et al., 2013). Other changes that occurred in recombinant carboxysomes involve the increased content of CsoS1 shell proteins, the reduced CsoS1D abundance, as well as the absence of CbbQO (the *cbbQ* and *cbbO* genes were not included in the expression construct) (Figure 3). All these structural alternations may collectively result in the higher size variation of recombinant α-carboxysomes and the discrepancy in the carbon-fixation performance between native and recombinant α-carboxysomes (Figure S1C-1E).

**Table 3.**
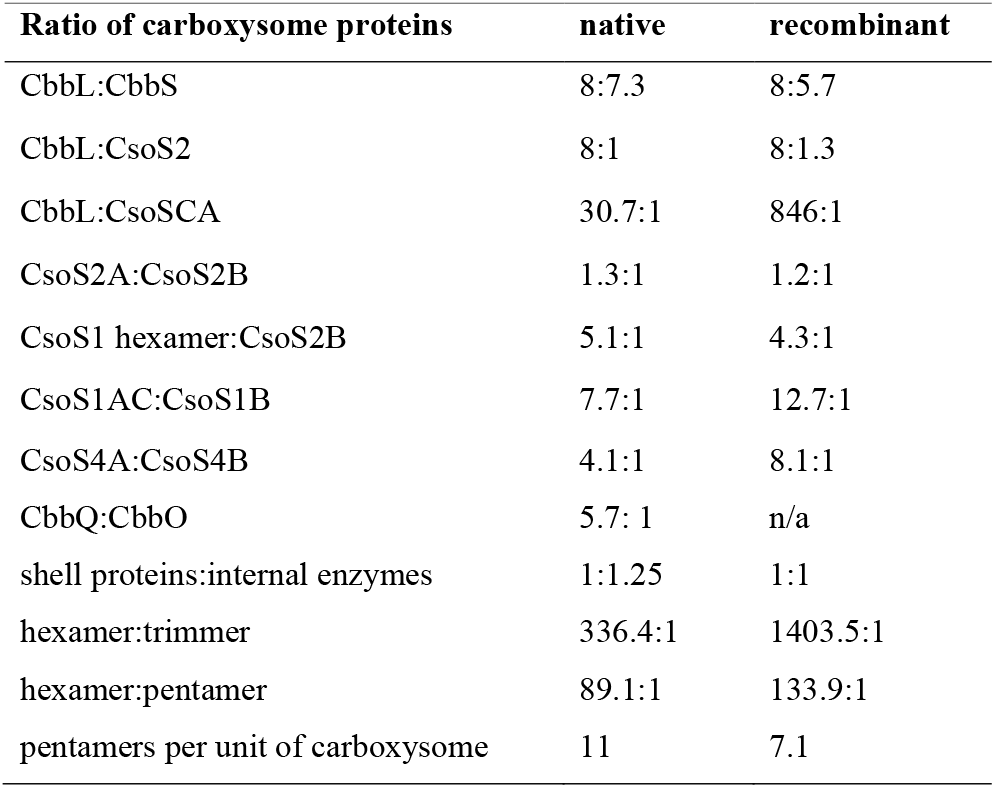
Stoichiometric ratios of protein components in α-carboxysomes. Interior proteins: CbbL, CbbS, CsoS2; Shell proteins: CsoS1, CsoS4.

CsoS2 in α-carboxysomes serve as the scaffolding protein that interlinks Rubisco and shells (Cai et al., 2015; Chaijarasphong et al., 2016; Rae et al., 2013). The CbbL:CsoS2 ratios in native and recombinant α-carboxysomes remain within a narrow range between 8:1 and 8:1.3 (Table 3), implicating the correlation between Rubisco and CsoS2, which is fundamental for Rubisco condensation and internal packing. Likewise, the CsoS2A:CsoS2B ratio remains relatively unaltered in native (ratio of 1.3:1) and recombinant (ratio of 1.2:1) α-carboxysomes.

### Organizational features of diverse carboxysomes

Peptide composition of the α-carboxysomes from the α-cyanobacterium *Prochlorococcus marinus* MED4 has been estimated based on standard protein gel profiles (Roberts et al., 2012). The *H. neapolitanus* α-carboxysomes (∼125 nm in diameter) are larger in diameter than the *Prochlorococcus* α-carboxysomes (∼90 nm in diameter). Consistently, the *H. neapolitanus* α-carboxysome has a 1.8-fold increased content of CsoS1 hexameric shell proteins (975 versus 539 copies) and encapsulates double copy numbers of CsoSCA proteins (58 versus 29) and nearly 3-fold more Rubisco enzymes (447 versus 152 copies). The experimentally determined Rubisco content fits well the theoretical estimate (411 copies for the *H. neapolitanus* carboxysome and 143 copies for the *Prochlorococcus* carboxysome), which were based on the carboxysome size and Kepler packing (Long et al., 2011; Whitehead et al., 2014). In contrast, Pdu microcompartments, with the diameter ranging from 90 to 130 nm, have a drastically higher shell:interior ratio (4.6:1) (Yang et al., 2020) than the *H. neapolitanus* α-carboxysome (0.8:1, Table 3), implying that Kepler packing of cargo enzymes is unlikely applicable to metabolosomes. The CsoSCA:CsoS1 ratio retain relatively constant in both native α-carboxysomes, presumably implicating their specific association within the carboxysomes. In contrast, CA in the Syn7942 β-carboxysomes, which encoded by the *ccaA* gene that is distant from the *ccm* operon, was demonstrated to have a varying abundance per carboxysome under different environmental conditions (Sun et al., 2019). It remains to be investigated if the CsoSCA content in α-carboxysomes is subject to environmental modulation.

A noteworthy feature of the *Prochlorococcus* α-carboxysome is that it contains only the full length of CsoS2 without the short isoform as the *H. neapolitanus* counterpart does, which might lead to formation of carboxysomes with reduced Rubisco loading capacity and overall size. However, the Rubisco:CsoS2 ratios in the α-carboxysomes from *H. neapolitanus* and *Prochlorococcus* remain relatively comparable (1:1 and 1:1.1, respectively), indicative of a general Rubisco encapsulation mechanism of α-carboxysomes. In the Syn7942 β-carboxysome, the ratio between Rubisco and the scaffolding protein CcmM varied in a range of 1:0.8 to 1:1.3, depending upon environmental conditions (Sun et al., 2019). Unlike the similar CsoS2A:CsoS2B ratios in native and recombinant α-carboxysomes, the CcmM35:CcmM58 ratios in the Syn7942 β-carboxysomes have a wide range of 1:1 to 11:1, and have been proved to be vital for carboxysome assembly (Long et al., 2011; Long et al., 2010).

Carboxysomes are highly modular structures with the capacity of incorporating foreign cargos, representing an ideal system in synthetic biology (Li et al., 2020). Advanced knowledge about the precise protein stoichiometry of functional carboxysome structures is essential for fine-tuning and reprogramming carboxysomes in native and heterogeneous organisms for metabolic enhancement and diverse biotechnological applications in new contexts (Liu et al., 2021). The QconCAT-based protein quantification technique could be broadly used in the studies of diverse BMC paralogs and protein organelles from their native origins and heterologous hosts.

## Methods

### Bacterial strains, growth conditions and carboxysome production

*H. neapolitanus* (*Halothiobacillus neapolitanus* Parker, Kelly and Wood ATCC 23641 C2) used in this work was acquired from ATCC (The American Type Culture Collection) as freeze-dried powder (Cannon et al., 2001; Hutchinson et al., 1965). Stock cells were maintained in liquid ATCC medium 290 (Hutchinson et al., 1967) or on ATCC 290 1.5% agar plates. Scale-up culture was grown similar to the protocol described previously (Dou et al., 2008), in the Vishniac and Santer medium (Vishniac and Santer, 1957) in a 5-liter fermenter (BioFlo 115, New Brunswick Scientific, USA) at 30°C. The pH of growth medium was maintained at 7.6 by constant supplement of 3 M KOH. Air supply was set at 500 L.min^-1^ for initial growth and reduced to 200 L.min^-1^ 24-48 hours prior to harvesting. Agitation was kept at 250-300 RPM. For expression of recombinant carboxysomes, the entire *cso* operon, as designed on *pHnCBS1D* reported previously (Bonacci et al., 2012), was fused on a pBAD33 arabinose inducible expression vector (Aigner and Wilson, 2017) using Gibson Assembly strategy (Gibson et al., 2009) with Gibson Assembly^®^ Master Mix from NEB. Primer sets used for assembly were listed in Table S5. For recombinant carboxysome expression in *E. coli*, seeding cultures containing chloramphenicol at a final concentration of 50 μg mL^-1^ were inoculated at 37°C in LB broth until reaching OD600 at 0.6, and then scaled up for induction with 1mM Arabinose at 20°C overnight.

### Carboxysome purification from *H. neapolitanus* and *E. coli*

The α-carboxysome purification from *H. neapolitanus* was modified from the protocol described previously (Heinhorst et al., 2006b). Sulfur-free Cells fractions were obtained by subsequential centrifugation at 12,000 g for 10 min, 300 g for 15 min and 12,000 g for 10 min in TEMB buffer (10 mM Tris-HCl, pH 8.0, 10 mM MgCl_2_, 20 mM NaHCO_3_, 1 mM EDTA). Cells in 15ml of TEMB were then incubated with egg lysosome for 1 hour at 30°C, and disrupted via glass beads beating (150-212 μm glass bead, acid washed, Sigma-Aldrich). The lysates were further treated with 33% (v/v) B-PERII (ThermoFisher Scientific, UK) and 0.5% (v/v) IGEPAL CA630 (Sigma-Aldrich) and placed on a rotary mixer for 2 hours. The unbroken cells and large membrane debris were removed by centrifugation at 9,000g for 10min. Crude CB enrichment was pelleted at 48,000 g for 30 min. The pellet was resuspended, briefly centrifuged at 9,000 g and then loaded to a step sucrose gradient (10% 20% 30% 35% 50% 60%) and ultra-centrifuged at 105,000 g for 35 min. The milky layer of enriched carboxysome was harvested at 35%-50% sucrose layers. Sucrose was removed by an additional round of ultracentrifuge after diluting with TEMB. The pure carboxysome pellet was resuspended in a small volume of TEMB. Unless indicated otherwise, all procedures were performed at 4°C. The carboxysome purification from *E. coli* was performed as described previously (Bonacci et al., 2012; So et al., 2004) with minor modifications. The step gradient of sucrose was kept the same as the one for native carboxysome isolation. Additionally, IGEPAL CA-630 detergent was used at 0.5% (v/v) after cell break to reduce membrane contaminants in final enrichments.

### SDS-PAGE analysis

SDS-PAGE were performed following standard procedures. 10 μg purified carboxysomal proteins or 100 μg whole cell fractions were loaded per-well on 15% polyacrylamide gels and stained with Coomassie Brilliant Blue G-250 (ThermoFisher Scientific, UK).

### Design, cell-free expression and purification of QconCAT standard

Absolute quantification for the carboxysomal proteins was designed by concatenated signature QconCAT peptides (Pratt et al., 2006) in a similar way to that described previously (Yang et al., 2020). In brief, up to three qualified peptide candidates, when available according to design principles (Pratt et al., 2006) were nominated to quantify each protein (Table S1). Candidate peptides were BLAST searched against protein database for both *H. neap* and *E. coli* to ensure their uniqueness. Due to the high level of sequence similarity of CsoS1A/B/C, CsoS2A/B and CsoS4A/B, peptides represent shared sequences as well as unique sequences were included (Figure S2). The DNA fragment encoding the above peptides, together with GluFib and cMyc (these peptides are used to quantify the standard) and 6x His-tag on N-terminal and C-terminal respectively were generated following ALACAT/Qbrick assembly strategy as reported previously (Johnson et al., 2021). The final DNA sequence (Table S2) was assembled into a pEU-E01 vector for cell-free expression using wheat germ lysate (CellFree Sciences Co., Ltd, Japan). Synthesis was completed with [^13^C_6_, ^15^N_4_] arginine and [^13^C_6_, ^15^N_2_] lysine (CK Isotopes Ltd, UK) using WEPR8240H full Expression kit following default protocols (2BScientific Ltd, UK). The QconCAT peptides were purified with Ni Sepharose suspension (GE Healthcare Ltd, UK) in centrifuge filters (Corning Costar Spin-X 0.45 um pore size cellulose acetate membrane, Merck, UK) following standard methods. The QconCAT was precipitated and resuspended in 30 μL 25 mM ammonium bicarbonate, with 0.1% (w/v) RapiGestTM SF surfactant (Waters, UK) and protease inhibitors (Roche cOmpleteTM, Mini, EDTA-free Protease Inhibitor Cocktail, Merck, UK).

### Proteomic analysis

The protein concentration of each sample was determined using a NanoDrop Spectrophotometer (ThermoFisher Scientific, UK). Protein (0.5 μg) was digested with 0.01 μg Trypsin Gold, Mass Spectrometry Grade (Promega, US) at 37°C overnight after pretreatment with 0.05% (w/v) RapiGestTM SF surfactant at 80°C for 10 mins, 4 mM dithiothreitol (Melford Laboratories Ltd., UK) at 60°C for 10 mins and 14 mM iodoacetamide at room temperature for 30 mins. Digestions were acidified with trifluoroacetic acid (Greyhound Chromatography and Allied Chemicals, UK) and then centrifuged at 13,000 g to remove insoluble, non-peptidic material. Four biological replicates for both native and recombinant carboxysomes were analyzed using an UltiMateTM 3000 RSLCnano system coupled to a Q Exactive™ HF Hybrid Quadrupole-Orbitrap™ Mass Spectrometer (ThermoFisher Scientific, UK) in data-dependent acquisition mode according to the protocol published (Johnson et al., 2021). The LC was operated under the control of Dionex Chromatography MS Link 2.14. The raw MS data files were loaded into Thermo Proteome Discoverer v.1.4 (ThermoFisher Scientific, UK) and searched against carboxysome QconCAT database using Mascot v.2.7 (Matrix Science London, UK) with trypsin as the specified enzyme. Each precursor ion was cleanly isolated using the high-resolution and high-scanning speed of the MS1 approach. A precursor mass tolerance of 10 ppm and a fragment ion mass tolerance of 0.01 Da were applied. Additionally, preparations of the four native and synthetic carboxysomes were analyzed by label-free quantification. Data analysis, including run alignment and peak picking, was carried out in Progenesis QI for Proteomics v4. The quantification data were also visualized and analyzed using Simplifi (simplify.protifi.com), which are available through permanent hyperlinks included in the text.

For single carboxysome quantitative normalization, relative quantifications from QconCAT were normalized both 12 pentamer coverage, and a single layer shell protein coverage of hexameric and pentameric proteins (Table S4). 12-pentamer normalization is done via assuming 60 copies of monomeric CsoS4A and CsoS4B in sum per carboxysome. For shell coverage normalization, the shell surface area is first calculated using TEM measured diameter with the following formula:

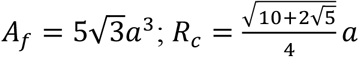

whereas *A*_*f*_ is total surface area, *α* is edge length, and *R*_*c*_ is the circumcized radius (refer to as the diameter). The hexameric counts were then calculated using the total surface area and diameters of CsoS1A hexamers in a layer as reported previously (Tsai et al., 2007).

### Electron Microscopy and data analysis

Electron microscopy was carried out as described previously (Faulkner et al., 2017). The purified carboxysomes (∼ 4 mg mL^-1^) were stained with 3% uranyl acetate on carbon grids and then inspected with FEI 120 kV Tecnai G2 Spirit BioTWIN TEM equipped with a Gatan Rio 16 camera. The diameters of carboxysomes were measured with ImageJ as described previously (Faulkner et al., 2017) and were statistically analyzed using Origin (OriginLab, Massachusetts, USA).

### Rubisco activity assays

Carbon fixation assay was carried out to determine carbon fixation capacities of purified native and recombinant carboxysomes as described previously using a 3-phosphoglycerate–dependent NADH oxidation coupled enzyme system (Bonacci et al., 2012). For both native and synthetic samples, four biological replicates that were isolated from different culture batches were assayed at 30°C, initiated with final concentrations of 0.0625 mM, 0.125 mM, 0.25 mM, 0.5 mM, 1 mM, and 2 mM of RuBP. The concentration of HCO_3-_ was set to 24 mM for all assays in this work.

## Acknowledgments

We thank Prof. Ian Prior and Mrs. Alison Beckett for the support of electron microscopy. This work was supported by the National Key R&D Program of China (2021YFA0909600 to L.-N.L.), the National Natural Science Foundation of China (32070109 to L.-N.L.), the Royal Society (URF\R\180030, RGF\EA\181061, RGF\EA\180233 to L.-N.L.), the Biotechnology and Biological Sciences Research Council Grant (BB/V009729/1, BB/M024202/1, BB/R003890/1 to L.-N.L. and BB/S020241/1 to R.J.B.), the Leverhulme Trust (RPG-2021-286 to L.-N.L.), and the International Postdoctoral Exchange Fellowship Program from China Postdoctoral Science Foundation (20180079 to T.C.).

## Author contributions

Y.Q., R.J.B. and L.-N.L. designed research; Y.Q., V.M.H., J.R.J., T.C., and G.F.D. performed research; Y.Q., V.M.H., Y.L., R.J.B. and L.-N.L. analyzed data; Y.Q. and L.-N.L. wrote the manuscript with contributions from other authors.

## Data availability

The mass spectrometry data are deposited to the public-accessible platform SimpliFi. The data for QconCAT quantification are available at: https://simplifi.protifi.com/#/p/dd32b950-2e5c-11eb-808d-871337eb317b. The data for label-free quantification search against *H. neapolitanus* protein database are available at: https://simplifi.protifi.com/#/p/e527f900-3e50-11eb-808d-871337eb317b. All other data are available from the corresponding author upon request.

## Competing Interests

The authors declare no conflict of interest.

## Supplemental Information

**Figure S1.**
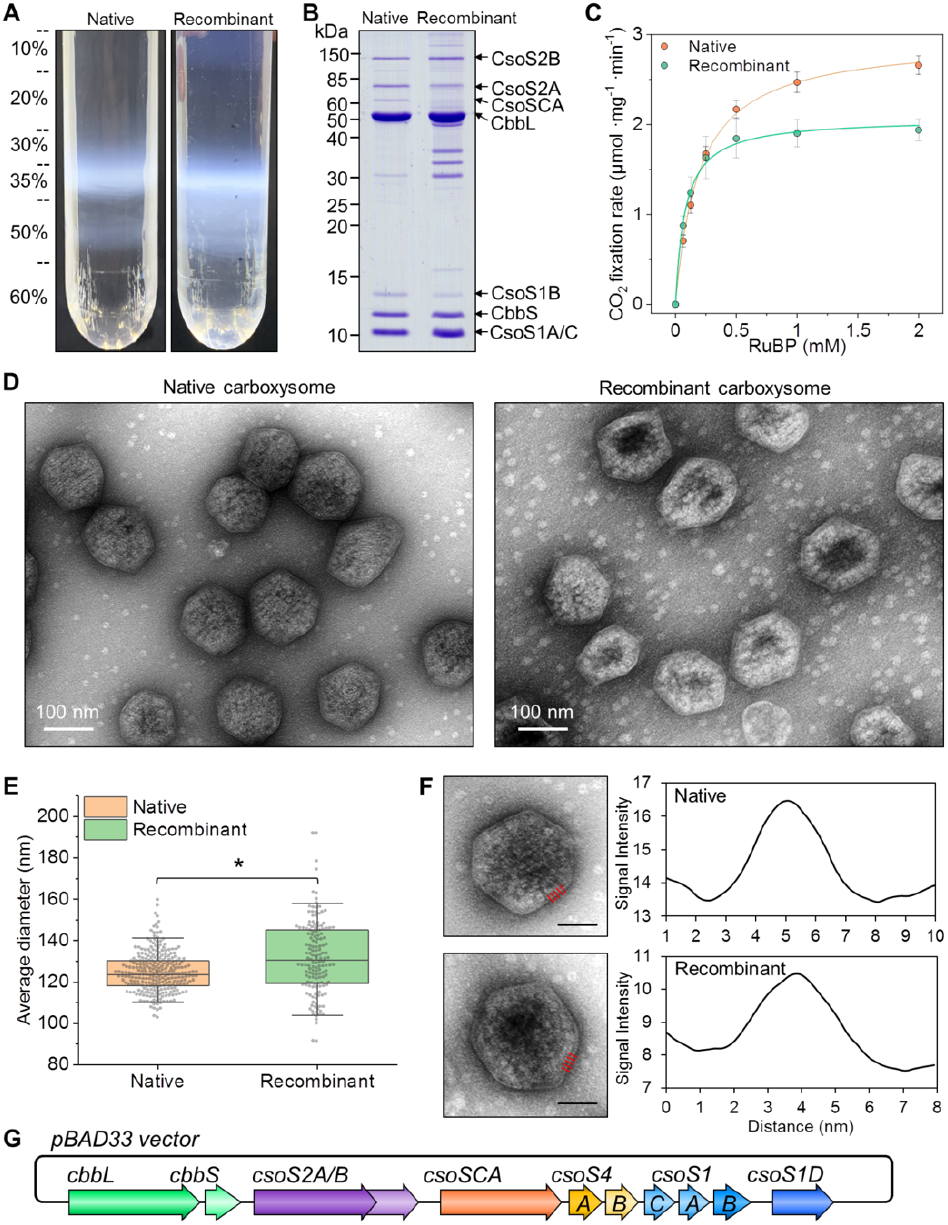
Purification and characterization of native and recombinant α-carboxysomes from *H. neapolitanus* and *E. coli*. **(A)** Sucrose gradient for carboxysome fractions. The milky band between 35%-50% sucrose fraction interface consists of enriched carboxysomes; **(B)** SDS-PAGE of purified carboxysomes isolated from four replicate batches of culture showing the majority bands of α-carboxysome proteins; **(C)** Carbon-fixation kinetics as a function of the RuBP concentrations revealed that native and recombinant carboxysomes possess *V*_max_ at 2.96 ± 0.09 and 2.07 ± 0.12 μmol.mg^-1.^min^-1^ and *K*_*m*(RuBP)_ at 0.20 ± 0.02 and 0.08 ± 0.02 mM, respectively. Data is shown as mean ± SD from four independent biological replicates; **(D)** TEM image of purified native and recombinant carboxysomes; (**E**) Boxplot distribution for diameters of purified native and recombinant carboxysomes, at 124.6 ± 9.6 nm (*n* = 272) and 131.8 ± 18.0 nm (*n* = 152), respectively. Significant difference of average diameter was confirmed with student t-test (*p* < 0.05); (F) Analysis on the shell thickness of native and recombinant α-carboxysomes. The shell thickness of native and recombinant α-carboxysomes is 5.3 ± 0.6 nm and 5.5 ± 0.8 nm, respectively (*n* = 100), implicating the single-layer shell architecture. The profile region for measurements were marked by red lines (Scale bar = 50 nm); **(G)** Recombinant α-carboxysome expression cassette containing *cso* operon and *cbbL/S*, plus BMC-T protein encoded gene csoS1D in the pBAD33 vector.

**Figure S2.**
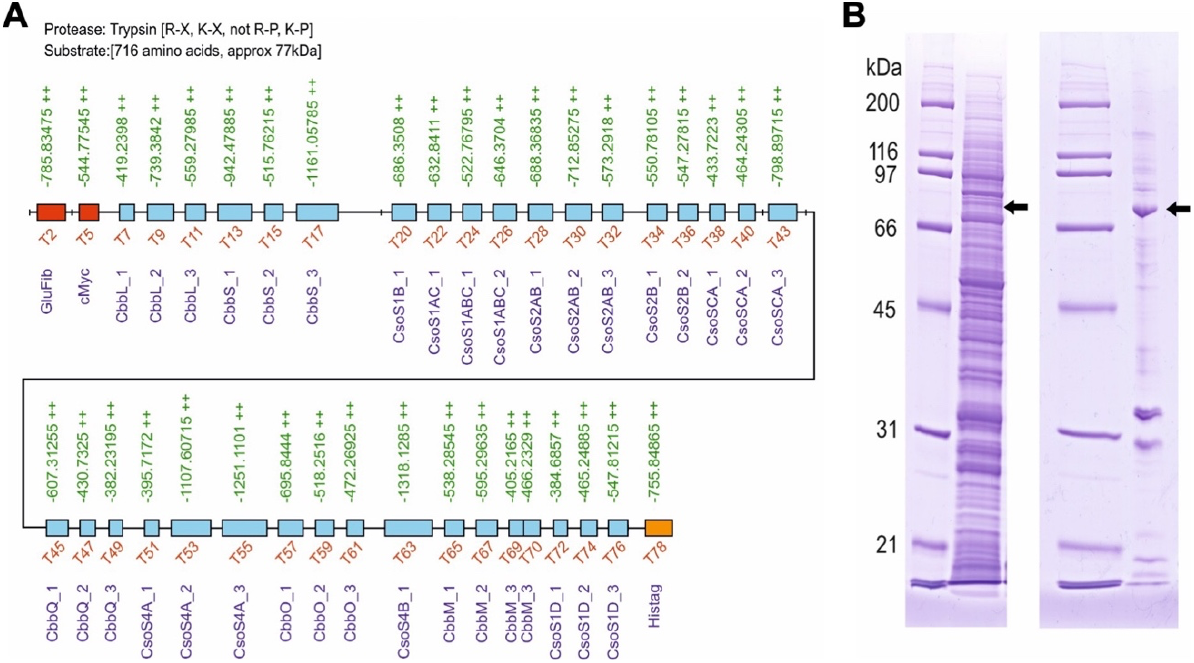
Structure and expression of the *H. neapolitanus* carboxysome QconCAT. (**A**) Schematic representation of the quantification concatamer, for quantification of proteins of interest. 35 peptides in QconCAT are represented by blue boxes. Values for the [M+2H^2+^] *m/z* peptide ions for the unlabeled QconCAT are aligned above each peptide (green text). The non-target quantification peptides (GluFib and cMyc) and the hexa-histidine tag for QconCAT purification are shaded in red and orange, respectively. (**B**) SDS-PAGE analysis of QconCAT expression and purification. The coding sequence of QconCAT peptide was sub-cloned into the cell-free expression vector pEU-E01-MCS (left). QconCAT was prepared by wheat germ cell-free synthesis in the presence of [^13^C_6_,^15^N_4_] arginine and [^13^C_6_,^15^N_2_] lysine and purified by virtue of the hexa-histidine tag (right). The QconCAT is denoted by the arrow.

**Figure S3.**
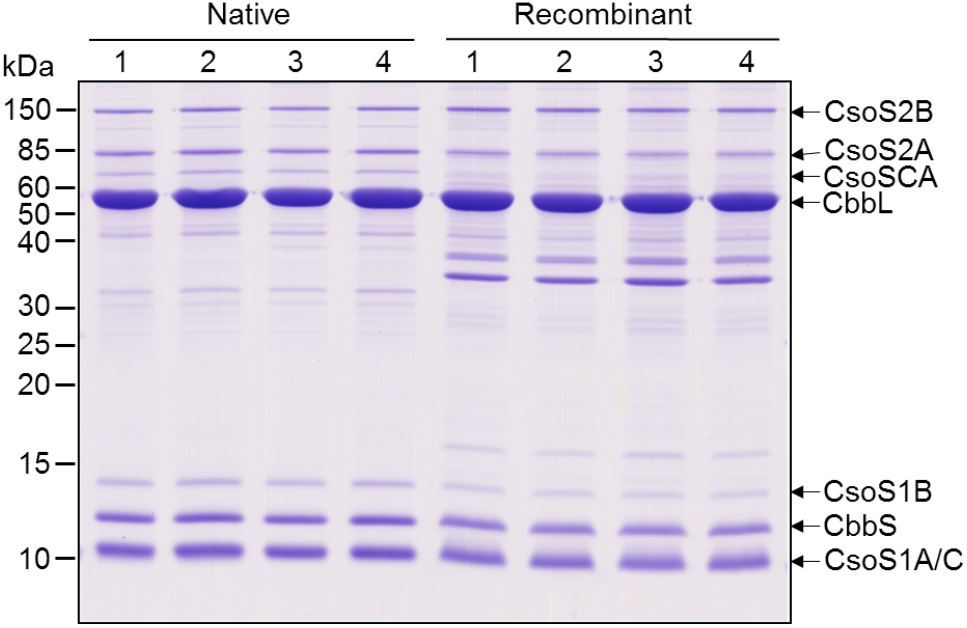
SDS-PAGE of purified carboxysomes from *H. neapolitanus* and *E. coli* with four biological replicates prepared for quantification by QconCAT mass spectrometry. 10 μg proteins are loaded per well.

**Figure S4.**
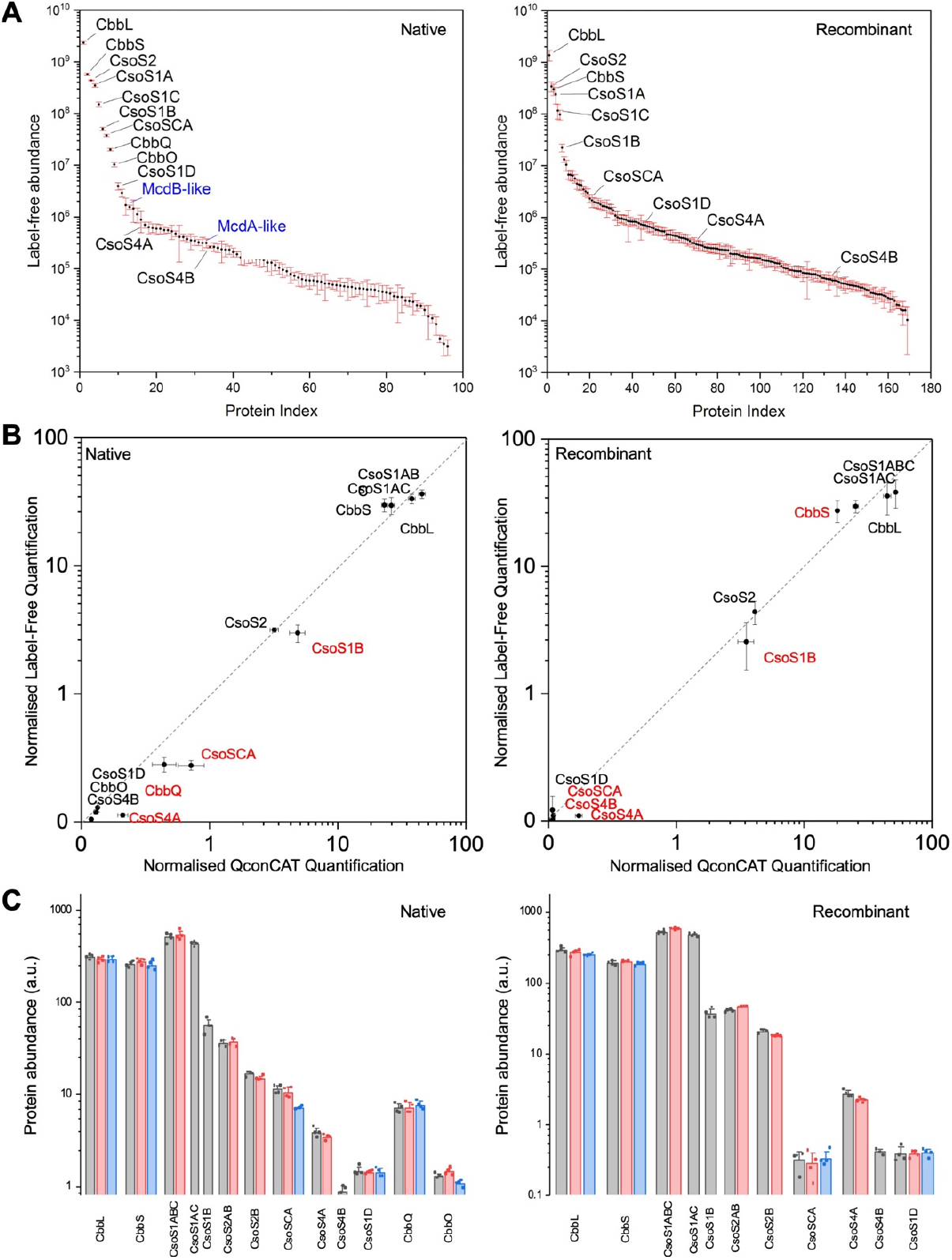
Evaluation of QconCAT and label-free quantification. (**A**) Protein index of all recognized proteins in label-free quantification. In native carboxysome samples, McdAB-like proteins are both detected. The full protein list provided in Supplemental File; (**B**) Comparison of QconCAT and label-free quantification. Quantification was normalized to equal total protein quantity. Proteins with the abundance difference greater than 30% from average are labelled in red; **(C)** Quantification of all QconCAT candidate peptides for *H. neapolitanus* carboxysomes. Overall good agreements are found within candidate peptides for same protein with the exception of peptide 3 of CsoSCA (34 ± 4 % lower than the other two) and CbbO (21 ± 3 %) in native carboxysome samples.

**Table S1.**
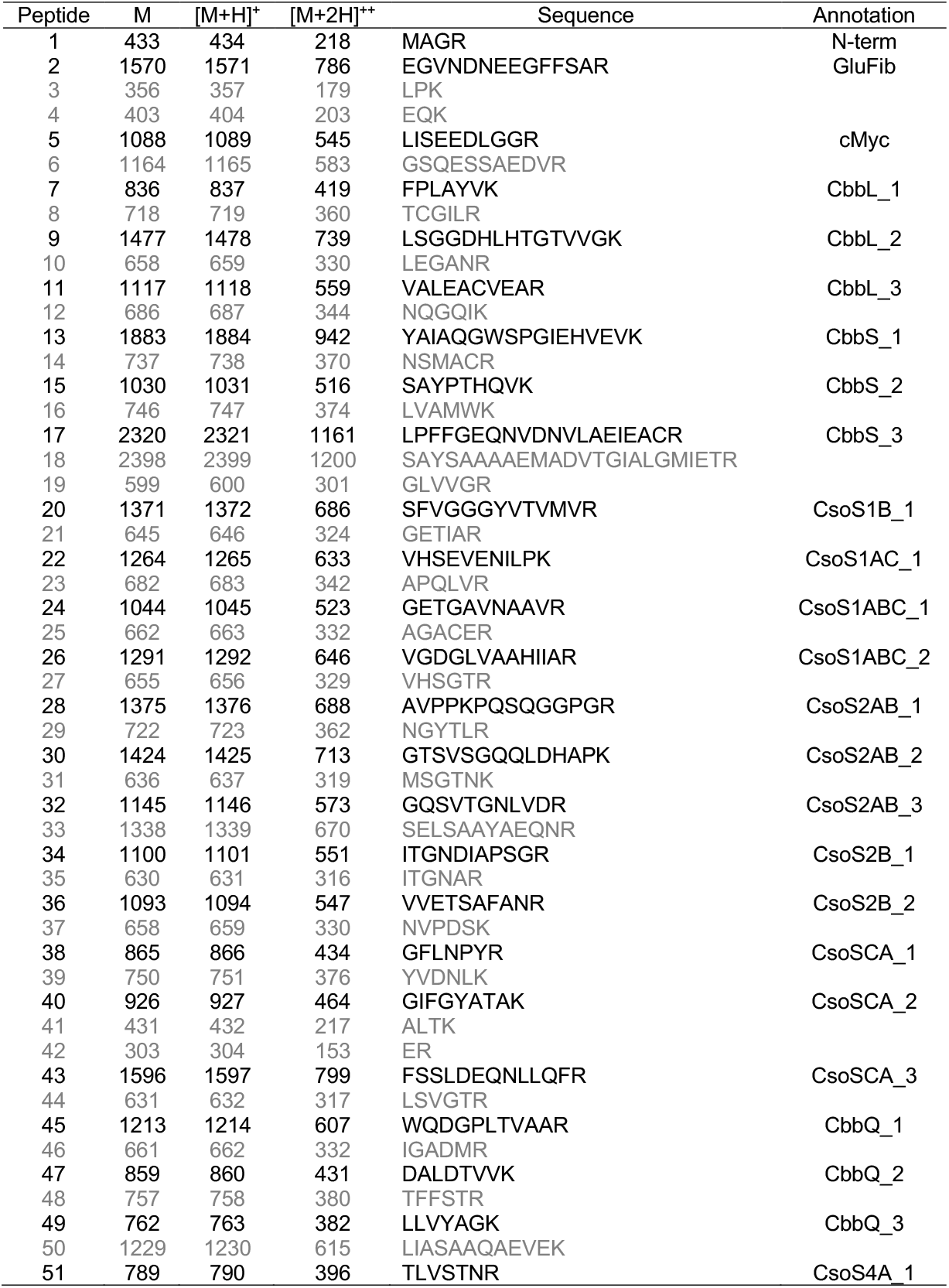
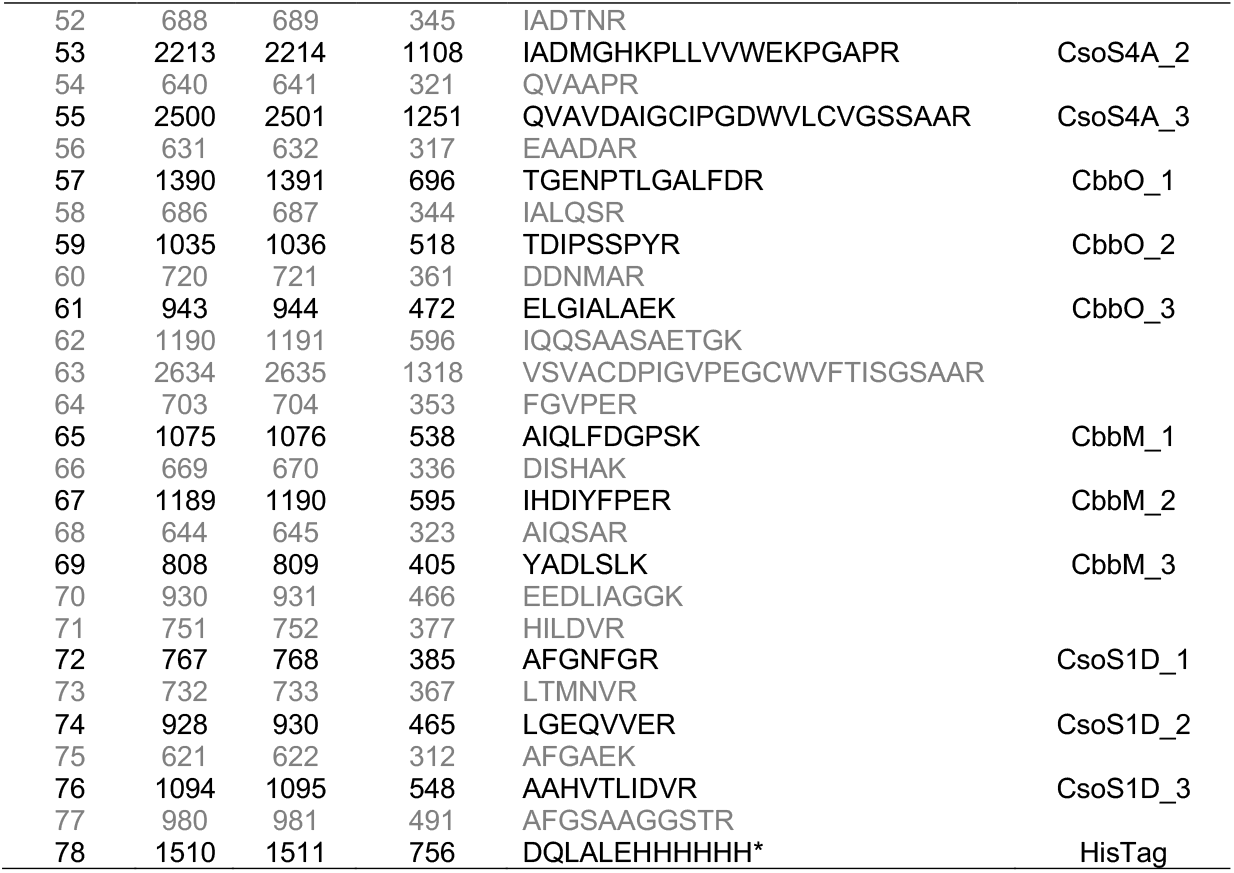
Peptides derived from tryptic proteolysis of the QconCAT for carboxysome protein quantification. The flanking sequences that recapitulate the true native primary sequence context, together with additional sequences that are derived from the loop assembly synthesis of the QconCAT are written in gray.

**Table S2.**
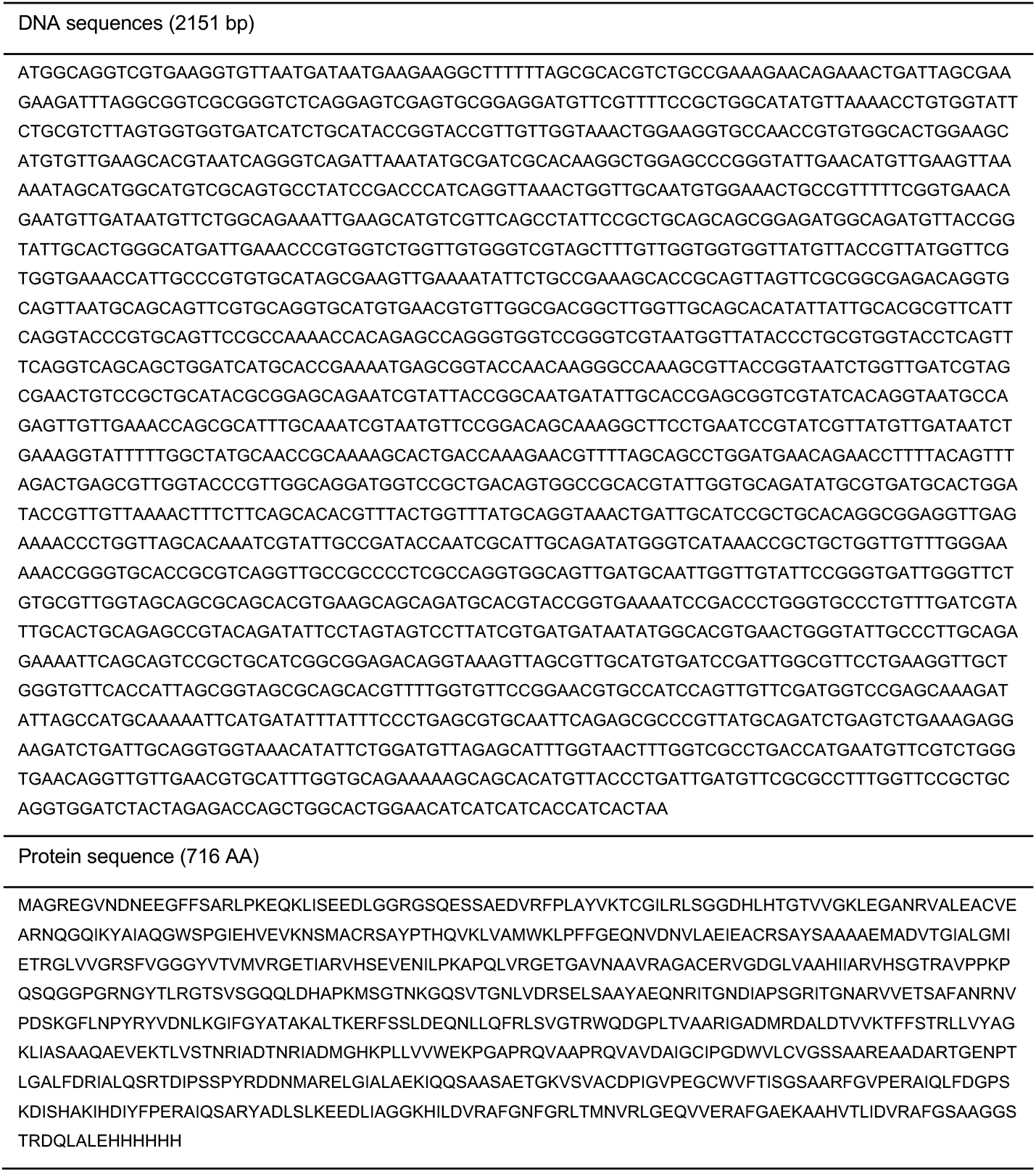
DNA and protein sequences of the recombinant carboxysome QconCAT peptide.

**Table S3.** Calculation of carboxysome surface area, shell hexamer content, and carboxysome diameter.

| | native α-carboxysome<br>(*n* = 272) | recombinant α-carboxysome<br>(*n* = 152) |
| --- | --- | --- |
| EM-measured diameter (nm) | 124.6 ± 9.6 | 131.8 ± 18.0 |
| Radius of circumscribed sphere (nm) | 62.3 ± 4.8 | 65.9 ± 9.0 |
| Facet side length (nm) | 65.5 ± 5.0 | 69.3 ± 9.4 |
| Carboxysome surface area (nm<sup>2</sup>) | 37407.3 ± 5864.6 | 42354.8 ± 11745.1 |
| CsoS1 hexamer width (nm)* | 6.64 | 6.64 |
| CsoS1 hexamer area (nm<sup>2</sup>)* | 38.9 | 38.9 |
| CsoS4 pentamer area (nm<sup>2</sup>)* | 30.3 | 30.3 |
| All facet hexamer counts | 977.6 ± 150.9 | 1080.2 ± 302.1 |
| Estimated Rubisco counts per carboxysome<sup>&</sup> | 410.7 ± 101.8 | 491.4 ± 216.1 |

\*CsoS1 and CsoS4 width/area obtained from previous publications (Tanaka et al., 2008; Tsai et al., 2007);
<sup>&</sup>Assumed packing densities of 74% (Kepler packing) (Whitehead et al., 2014) in the proposed carboxysome model based on the measured carboxysome diameters.

**Table S4.** Absolute protein abundance per native and recombinant carboxysome based on 12-pentamer occupation and surface area coverage.

| Protein | native α-carboxysome | | recombinant α-carboxysome | |
| --- | --- | --- | --- | --- |
|  | 12 pentamers | CsoS1 coverage* | 12 pentamers | CsoS1 coverage* |
| CbbL | 490.2 ± 20.7 | 447.0 ± 18.9 | 717.0 ± 53.1 | 426.0 ± 31.5 |
| CbbS | 433.4 ± 20.7 | 395.1 ± 18.8 | 515.3 ± 20.2 | 306.2 ± 12 |
| CsoS1AC | 945.9 ± 69.1 | **862.5 ± 63** | 1683.9 ± 75.2 | **1000.5 ± 44.7** |
| CsoS1B | 123.1 ± 27.1 | **112.2 ± 24.7** | 132.7 ± 15.9 | **78.8 ± 9.4** |
| CsoS2A | 271.7 ± 51.8 | 247.7 ± 47.3 | 513.3 ± 15.7 | 305 ± 9.3 |
| CsoS2B | 210 ± 16.4 | 191.5 ± 15 | 419.1 ± 21.3 | 249 ± 12.6 |
| CsoSCA | 63.9 ± 3.9 | 58.3 ± 3.6 | 3.4 ± 0.9 | 2.0 ± 0.5 |
| CsoS4A | **9.6 ± 0.4** | 8.8 ± 0.3 | **10.7 ± 1.2** | 6.3 ± 0.7 |
| CsoS4B | **2.4 ± 0.4** | 2.2 ± 0.3 | **1.3 ± 0.9** | 0.8 ± 0.5 |
| CsoS1D | 3.2 ± 0.2 | **2.9 ± 0.2** | 1.4 ± 0.1 | **0.8 ± 0.1** |
| CbbQ | 16 ± 1 | 14.6 ± 0.9 | N/A | N/A |
| CbbO | 16.9 ± 1.4 | 15.4 ± 1.3 | N/A | N/A |

\*Based on surface area of ideal icosahedral with diameters measured from EM, 124.6 ± 9.6 nm ( $n$ = 272) and 131.8 ± 18.0 nm ( $n$ = 152) for native and recombinant carboxysomes, respectively. Value used for standardization was displayed in bold. Calculation was described in Methods.

**Table S5.**
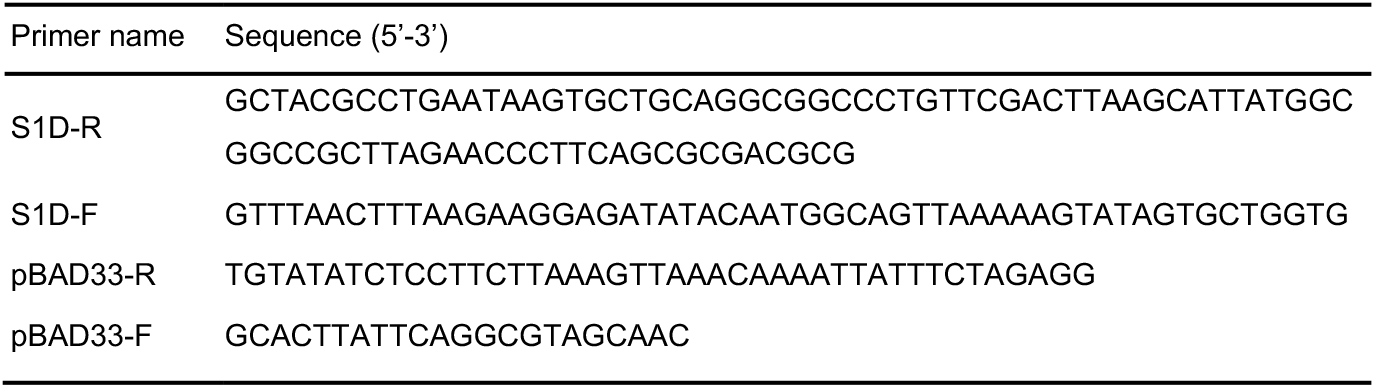
Primer sets used for pBAD33-CBS1D construction.

